# TRA2A-induced upregulation of LINC00662 regulates blood-brain barrier permeability by affecting ELK4 mRNA stability in Alzheimer’s microenvironment

**DOI:** 10.1101/848408

**Authors:** Qianshuo Liu, Lu Zhu, Xiaobai Liu, Jian Zheng, Yunhui Liu, Xuelei Ruan, Shuo Cao, Heng Cai, Zhen Li, Yixue Xue

## Abstract

The blood-brain barrier (BBB) has an important significance in maintenance and regulation of the neural microenvironment. The occurrence of BBB disruption is the pathological change of early Alzheimer’s disease (AD). RNA-binding proteins and long non-coding RNAs are closely related to the regulation of BBB permeability. Our study was performed to demonstrate TRA2A/LINC00662/ELK4 axis that regulates BBB permeability in AD microenvironment. In Aβ_1-42_-incubated microvascular endothelial cells (ECs) of BBB model *in vitro*, TRA2A and LINC00662 were enriched. TRA2A increased the stability of LINC00662 by binding with it. The knockdown of either TRA2A or LINC00662 decreased the BBB permeability *via* upregulating the levels of tight junction-related proteins. ELK4 was downregulated in BBB model *in vitro* in AD microenvironment. LINC00662 mediated the degradation of ELK4 mRNA by SMD pathway. The downregulated ELK4 increased the permeability of BTB by inducing the tight junction-related proteins. TRA2A/LINC00662/ELK4 axis is important in the regulation of BBB permeability in AD microenvironment, which would be a new molecular target for AD treatment.

## Introduction

Alzheimer’s disease (AD), as the most common cause of dementia, seriously affects human health and quality of life^[1]^. At present, since the pathogenesis of AD remains unclear, the existing treatment programs are not effective^[2]^. The occurrence of blood-brain barrier (BBB) disruption is the pathological change of early AD. The increase of BBB permeability aggravates the degeneration of blood vessels and neurons^[3, 4]^. Therefore, function changes of BBB in AD have been a great challenge for identification of pathogenesis and treatments^[5]^. BBB is composed of cerebral microvascular endothelial cells (ECs), pericytes, extracellular matrix and podocytes of perivascular astrocytes^[6]^. The tight junctions between adjacent ECs play a critical role in maintaining BBB integrity. BBB disruption in AD mainly reduces Aβ clearance^[4]^, promotes inflow into the brain of neurotoxic substances, and is associated with inflammatory and immune responses, which can initiate multiple pathways of neurodegeneration^[7]^.

RNA-binding proteins (RBPs) are a class of proteins that bind to specific RNA. RBPs can influence the transcription process from alternative splicing, polyadenylation, and nuclear export to cytoplasmic localization, stability, and translation. It has been reported that RBPs, such as Embryonic lethal abnormal vision like protein family (ELVAL), TAR DNA-binding protein 43 (TDP43), and eukaryotic initiation factor 2 (eIF2α), play pivotal regulatory roles in the development of neurodegenerative diseases including AD^[8–10]^. TRA2A is located at 7p15.3 and significantly impacts the splicing regulatory process of pre-mRNA^[11]^. TRA2A is highly expressed in the neurodegenerative disease, fragile X-associated tremor/ataxia syndrome (FXTAS). TRA2A aggregates in FXTAS mouse models and in post-mortem human samples, which indicates TRA2A is associated with FXTAS occurrence^[12]^. TRA2A regulates alternative splicing in triple-negative breast cancers, which result in chemotherapy drugs resistance and tumor progression^[13]^. TRA2A is upregulated in glioma and promotes malignant biological behavior of glioma^[14]^. Additionally, RBPs mediated regulation of vascular function is involved with long non-coding RNAs (lncRNAs)^[15]^.

LncRNAs are a group of non-coding RNA transcripts comprising more than 200 nucleotides and lacking apparent open reading frames. It is reproted that lncRNAs are significant in biological processes on the transcriptional and post-transcriptional levels, regulating gene transcription, pre-mRNA processing, mRNA stability, protein translation, *etc*. LncRNAs may provide novel approaches for early diagnosis and treatments of AD^[16]^. LINC00662 is located at 19q11 and relates to tumorigenesis^[17, 18]^. However, the regulatory role and potential mechanisms of RBPs and lncRNAs affecting the function of ECs should be further elucidated.

ETS-domain protein 4 (ELK4) (also known as SAP1) is a member of the Ets transcription factor family. As a cofactor of serum inflammatory factors, ELK4 participates in inflammatory response^[19]^, insulin dependence regulation^[20]^, *etc*. In gliomas, down-regulation of ELK4 can decrease the expression of Mcl-1 and induce the sensitivity of tumor cells to apoptosis^[21]^. ELK4 as a transcriptional regulator participates in the expression of neuronal plasticity regulatory genes under the condition of sleep deprivation^[22]^. By searching the bioinformatics database RepeatMasker, we found that there is a Alu element in ELK4’s 3’-UTR region. We also predicted that the putative binding site exists between LINC00662 and ELK4 3’-UTR region using bioinformatics software IntaRNA.

STAU1-mediated mRNA decay (SMD) pathway is an mRNA degradation pathway that regulates biological processes. Stau1 binding sites (SBSs) can be formed by base-pairing between a Alu element within lncRNA and another Alu element within the 3’UTR of a target mRNA. SMD regulates the expressions of target mRNAs that harbor SBSs by recruting STAU1 and UPF1^[23]^. SMD pathway is involved with tumorigenesis^[15]^ and cell differentiation^[24]^. It is reported that STAU1 is increased in Spinocerebellar ataxia type 2 (SCA2), a neurodegenerative disease model, and involved in abnormal RNA metabolism^[25]^. Thus, we inferred that SMD pathway may happen in neurodegenerative diseases.

In present study, endogenous expressions of TRA2A, LINC00662 and ELK4 in ECs after Aβ_1-42_ incubation were detected, and interactions among them were analyzed to confirm the regulatory mechanisms for BBB permeability. Our study may provide a new target for AD treatment regard of BBB.

## Results

### TRA2A was highly expressed in Aβ_1-42_-incubated ECs, and knockdown of TRA2A attenuatd BBB permeability in AD microenvironment

In this study, Aβ_1-42_-incubated human brain microvascular endothelial cells (hCMEC/D3) were used to simulate the model of BBB in AD microenvironment. To verify the role of TRA2A in BBB permeability in AD microenvironment, TRA2A levels in Aβ_1-42_-incubated ECs was detected by qRT-PCR and Western blot. As shown in Figure. 1-A,B, TRA2A mRNA and protein levels in Aβ_1-42_-incubated ECs were remarkablely increased. TRA2A was knocked down to futher investigated its function. When the BBB model *in vitro* was constructed using ECs transfected with shTRA2A plasmid in AD microenvironment, TEER values, HRP flux were detected to analyze the integrity and permeability of BBB, respectively. Compared with shNC group, TEER in shTRA2A group increased (Figure. 1-C), HRP flux attenuated significantly (Figure.1-D), indicating that TRA2A konckdown decreased BBB permeability in AD microenvironment. The expression levels and distribution of ZO-1, occludin and claudin-5 were detected to clarify the possible mechanisms of BBB permeability. As shown in Figure. 1-E,F, knockdown of TRA2A significantly induced ZO-1, occludin and claudin-5 mRNA and protein expression levels in Aβ_1-42_-incubated ECs. Moreover, consistent with the western blot results, immunofluorescence showed that TRA2A knockdown increased the expression of ZO-1, occludin and claudin-5, with discontinuous distribution to relatively continuous distribution on the boundaries of Aβ_1-42_-incubated ECs.

**Figure 1.**
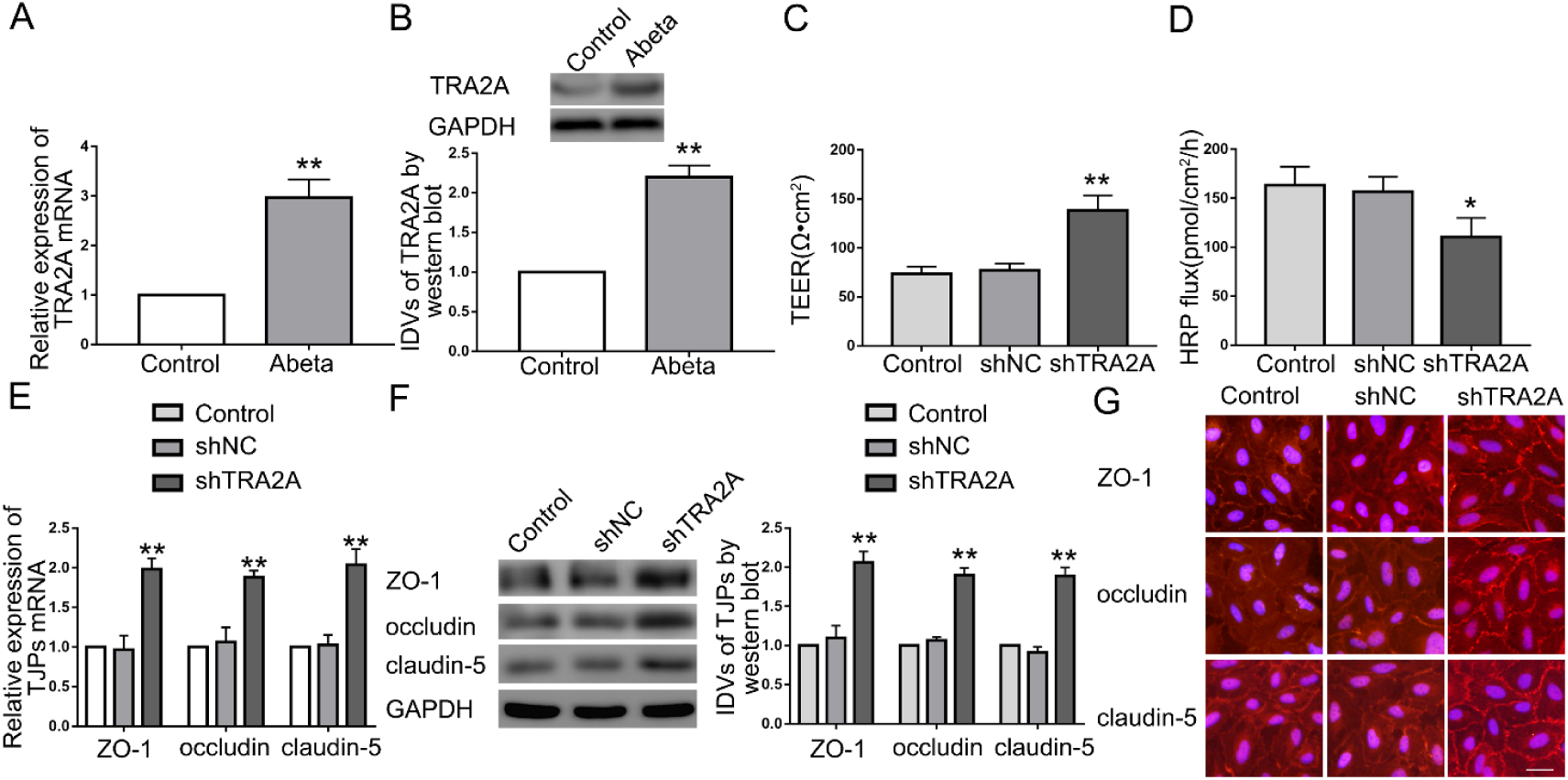
TRA2A expression in Aβ_1-42_-incubated ECs and TRA2A regulated BBB permeability in AD microenvironment. (A) Relative mRNA expression of TRA2A in ECs pre-incubated with Abeta_1-42_ by qRT-PCR. Data are presented as mean ± SD (n = 3, each). *\*\*P* < 0.01 *versus* control group. (B) Relative TRA2A protein levels in ECs pre-incubated with Abeta_1-42_ by western blot. *\*\*P* < 0.01 *versus* control group. (C and D) Effects of TRA2A on TEER values (C) and HRP flux (D). Data are presented as mean ± SD (n = 3, each). *\*P* < 0.05 *versus* shNC group. *\*\*P* < 0.01 *versus* shNC group. (E) Effects of TRA2A on ZO-1, occludin, and claudin-5 expression levels determined by qRT-PCR. Data are presented as mean ± SD (n = 3, each). *\*\*P* < 0.01 *versus* shNC group. (F) Effects of TRA2A on ZO-1, occludin, and claudin-5 expression levels determined by western blot. Data are presented as mean ± SD (n = 3, each). *\*\*P* < 0.01 *versus* shNC group. (G) Effects of TRA2A on ZO-1, occludin, and claudin-5 expression levels and distribution determined by immunofluorescence staining (n = 3, each). ZO-1, occludin, and claudin-5 (red) were labeled with secondary antibody against anti-ZO-1, anti-occludin, and anti-claudin-5 antibody, respectively, and nuclei (blue) were labeled with DAPI. Scale bar represents 30 μm.

### LINC00662 was up-regulated in Aβ_1-42_-incubated ECs, and knockdown of LINC00662 decreased BBB permeability in AD microenvironment

To investigate lncRNAs that involved in TRA2A-mediated regulation on BBB permeability, we used lncRNA microarray. It was found LINC00094, LINC00662 and FIRRE were significantly downregulated in ECs transfected with shTRA2A plasmid (Figure 2-A). Further, the lncRNAs level in shTRA2A ECs was investigated by qRT-PCR. As Figure 2-B shows, LINC00662 expression was remarkablely attenuated in Aβ_1-42_-incubated ECs treated with shTRA2A. Moreover, LINC00662 was upregulated in Aβ_1-42_-incubated ECs (Figure 2-C). TEER value in shLINC00662 group increased significantly (Figure. 2-D), HRP flux attenuated significantly (Figure. 2-E) compared with shNC group. The mechanistic studies showed, the mRNA and protein levels of ZO-1, occludin, and claudin-5 in Aβ_1-42_-incubated ECs were promoted after LINC00662 knockdown (Figure. 2-F, G). Immunofluorescence confirmed that downregulation of LINC00662 induces ZO-1, occludin and claudin-5, with more continuous distribution on the boundaries of ECs (Figure. 2-H).

**Figure 2.**
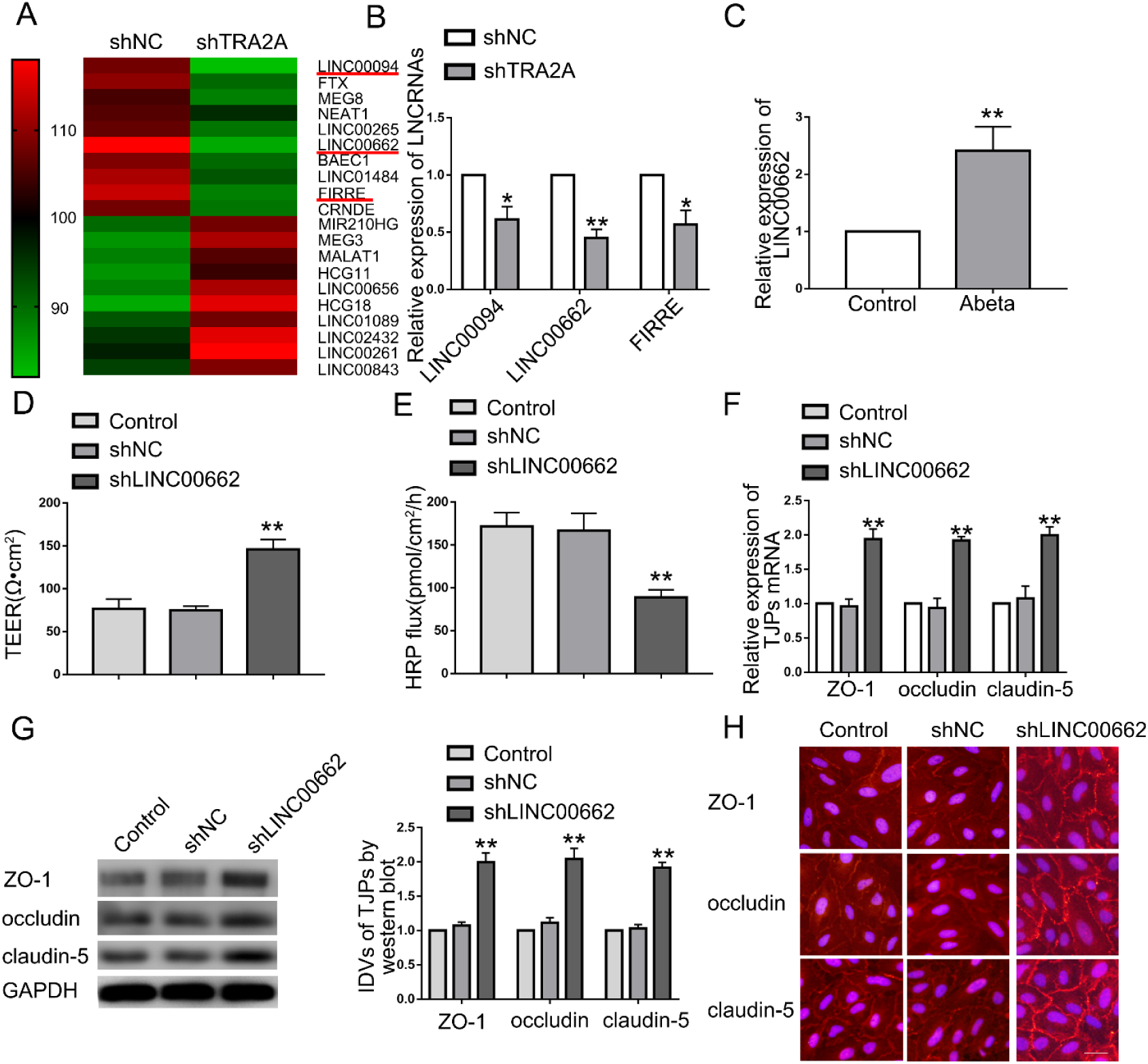
LINC00662 expression in Aβ_1-42_-incubated ECs and LINC00662 regulated BBB permeability in AD microenvironment. (A) LncRNA microarray analysis was performed in ECs treated with shTRA2A. Red indicates high relative expression and green indicates low relative expression. (B) Relative expression levels of LINC00094, LINC00662, and FIRRE determined by qRT-PCR. Data represent mean ± SD (n = 3, each). *\*P* < 0.05 *versus* shNC group. *\*\*P* < 0.01 *versus* shNC group. (C) Relative expression of LINC00662 in ECs pre-incubated with Aβ_1-42_ by qRT-PCR. Data are presented as mean ± SD (n = 3, each). *\*P* < 0.05 *versus* control group. (D and E) Effects of LINC00662 on TEER values (D) and HRP flux (E). Data are presented as mean ± SD (n = 3, each). *\*P* < 0.01 *versus* shNC group. (F) Effects of LINC00662 on ZO-1, occludin, and claudin-5 expression levels determined by qRT-PCR. Data are presented as mean ± SD (n = 3, each). *\*P* < 0.01 *versus* shNC group. (G) Effects of LINC00662 on ZO-1, occludin, and claudin-5 expression levels determined by western blot. Data are presented as mean ± SD (n = 3, each). *\*P* < 0.01 *versus* shNC group. (H) Effects of LINC00662 on ZO-1, occludin, and claudin-5 expression levels and distribution determined by immunofluorescence staining (n = 3, each). ZO-1, occludin, and claudin-5 (red) were labeled with secondary antibody against anti-ZO-1, anti-occludin, and anti-claudin-5 antibody, respectively, and nuclei (blue) were labeled with DAPI. Scale bar represents 30 μm.

### LINC00662 was involved in TRA2A-mediated regulation of BBB permeability

As shown in Figure 3-A, the expression level of LINC00662 in shTRA2A group was significantly lower than that in shNC group using qRT-PCR. To explore the correlation between TRA2A and LINC00662, RIP assays and RNA pull-down assays were used to confirm that TRA2A binds to LINC00662 directly. As shown in Figure. 3-B, the enrichment of LINC00662 in Anti-TRA2A group was higher than anti-IgG group. The level of TRA2A captured by Anti-LINC00662 group was significantly higher than that in anti-antisense group (Figure. 3-C). Moreover, qRT-PCR demonstrated that novel LINC00662 level in shNC group and shTRA2A group is not statistically significant (Figure. 3-D). As shown in Figure. 3-E, the half-life of LINC00662 in shTRA2A group was significantly impaired than that in shNC group. LINC00662 knockdown enhanced the increase in TEER values (Figure. 3-F) and the impairment in HRP flux (Figure. 3-G) caused by TRA2A knockdown. LINC00662 knockdown magnified the increase in ZO-1, occludin, claudin-5 expression caused by TRA2A knockdown (Figure. 3-H). These results indicated that TRA2A increased BBB permeability by stabilizing LINC00662.

**Figure 3.**
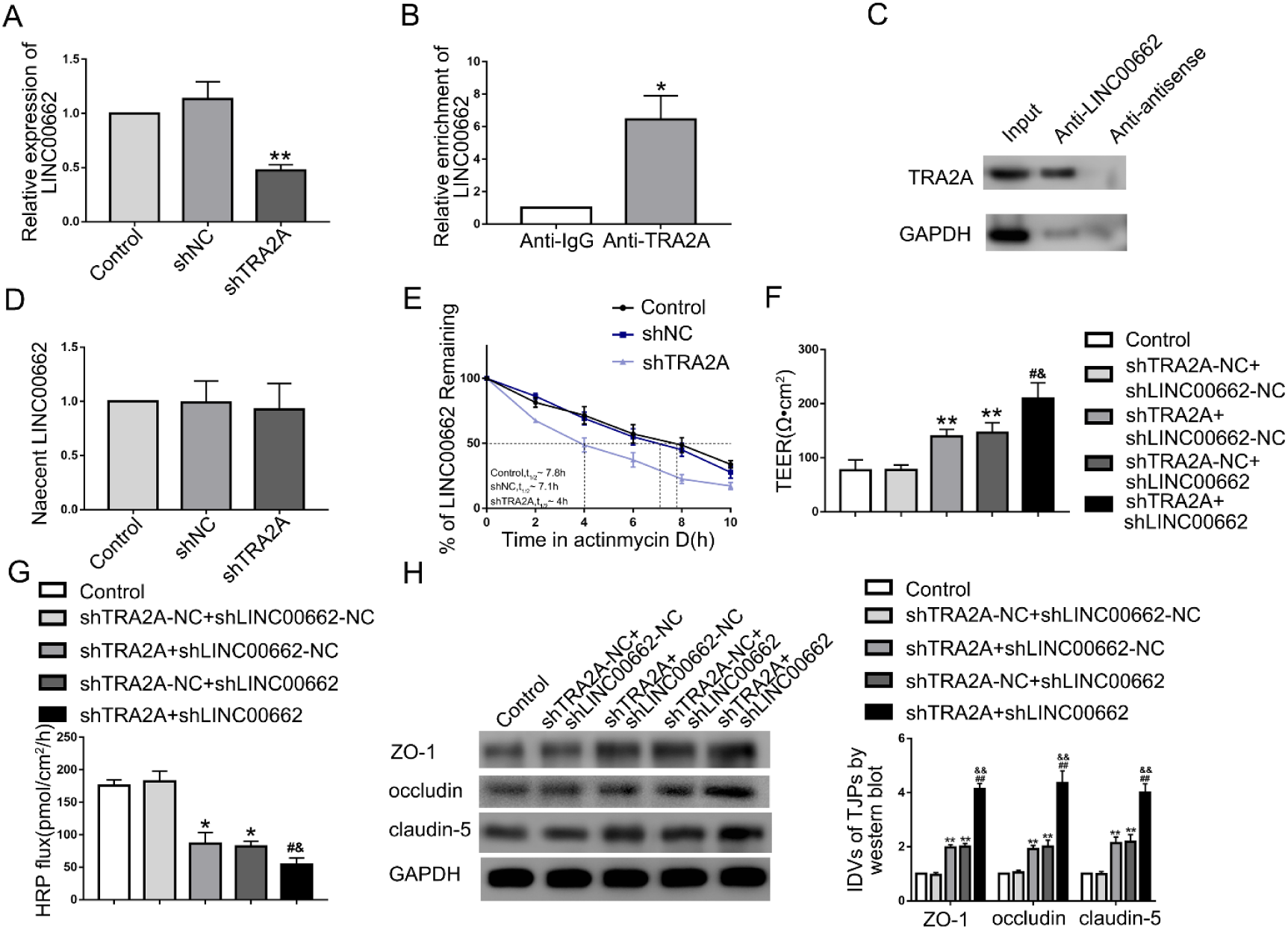
TRA2A konckdown decreased BBB permeability by stabilizing LINC00662. (A) Effects of TRA2A on LINC00662 expression determined by qRT-PCR. Data are presented as mean ± SD (n = 4, each). *\*\*P* < 0.01 *versus* shNC group. (B) RNA-IP confirmed the binding interaction between TRA2A and LINC00662. Relative enrichment was measured by qRT-PCR; Data are presented as mean ± SD (n = 3, each). *\*\*P* < 0.01 *versus* Anti-IgG group. (C) RNA pull-down indicated that TRA2A bind to LINC00662 directly. (D) The graph shows nascent LINC00662 in Aβ_1-42_-incubated ECs; data are presented as mean ± SD (n = 3, each). (E) The graph shows LINC00662 levels at different times treated by ActD in the control group, shNC group, and shTRA2A group. (F and G) Effects of TRA2A and LINC00662 knockdown on TEER values (F) and HRP flux (G). Data are presented as mean ± SD (n = 3, each). *\*\*P* < 0.01 *versus* shTRA2A-NC + shLINC00662-NC group, ^#^*P* < 0.05 *versus* shTRA2A + shLINC00662-NC, *^&^P* < 0.05 *versus* shTRA2A-NC + shLINC00662 group. (H) Effects of TRA2A and LINC00662 knockdown on ZO-1, occludin, and claudin-5 expression levels determined by western blot. Data are presented as mean ± SD (n = 3, each). *\*\*P* < 0.01 *versus* shTRA2A-NC + shLINC00662-NC group, ^##^*P* < 0.01 *versus* shTRA2A + shLINC00662-NC, *^&&^P* < 0.01 *versus* shTRA2A-NC + shLINC00662 group.

### ELK4 was down-regulated in Aβ_1-42_-incubated ECs, and attenuated BBB permeability by regulating ZO-1, occludin, claudin-5

When LINC00662 was knockdown in Aβ_1-42_-incubated ECs, microarray analysis showed ZEB1, ELK4 and ART1 are three most abundant (Figure. 4-A). The increase of the ELK4 expression investigated by qRT-PCR was the most significant (Figure. 4-B). ELK4 protein levels was highly expressed in shLINC00662 group (Figure 4-C). As shown in Figure. 4-D,E, the mRNA and protein expression levels of ELK4 in Aβ_1-42_-incubated ECs was lower than that in control group. To verify the role of ELK4 in BBB permeability in AD microenvironment, we knocked down and overexpressed ELK4 to elucidate its function. As shown in Figure. 4-F,G, compared with ELK4-NC group, TEER in ELK4 group increased significantly, HRP flux decreased significantly; compared with shELK4-NC group, TEER in shELK4 group decreased significantly and HRP flux increased significantly. Moreover, ZO-1, occludin, claudin-5 mRNA and protein exhibited higher expression level in ELK4 group, and ELK4 knockdown inhibited ZO-1, occludin and claudin-5 expression (Figure. 4-H,I). Subsequent immunofluorescence showed that ZO-1, occludin and claudin-5 were upregulated in ELK4 group, which exhibited relative continuous distribution on the boundaries of Aβ_1-42_-incubated ECs. ELK4 knockdown leaded to the opposite result (Figure. 4J). We further explored the role of TRA2A and LINC00662 on the expression of ELK4. Compared with shTRA2A-NC+shLINC00662-NC group, shTRA2A+shLINC00662-NC group and shTRA2A-NC+shLINC00662 group exhibited higher expression of ELK4. The expression of ELK4 in shTRA2A+shLINC00662 group was significantly increased compared with shTRA2A+shLINC00662-NC group and shTRA2A-NC+ shLINC00662 group respectively.

**Figure 4.**
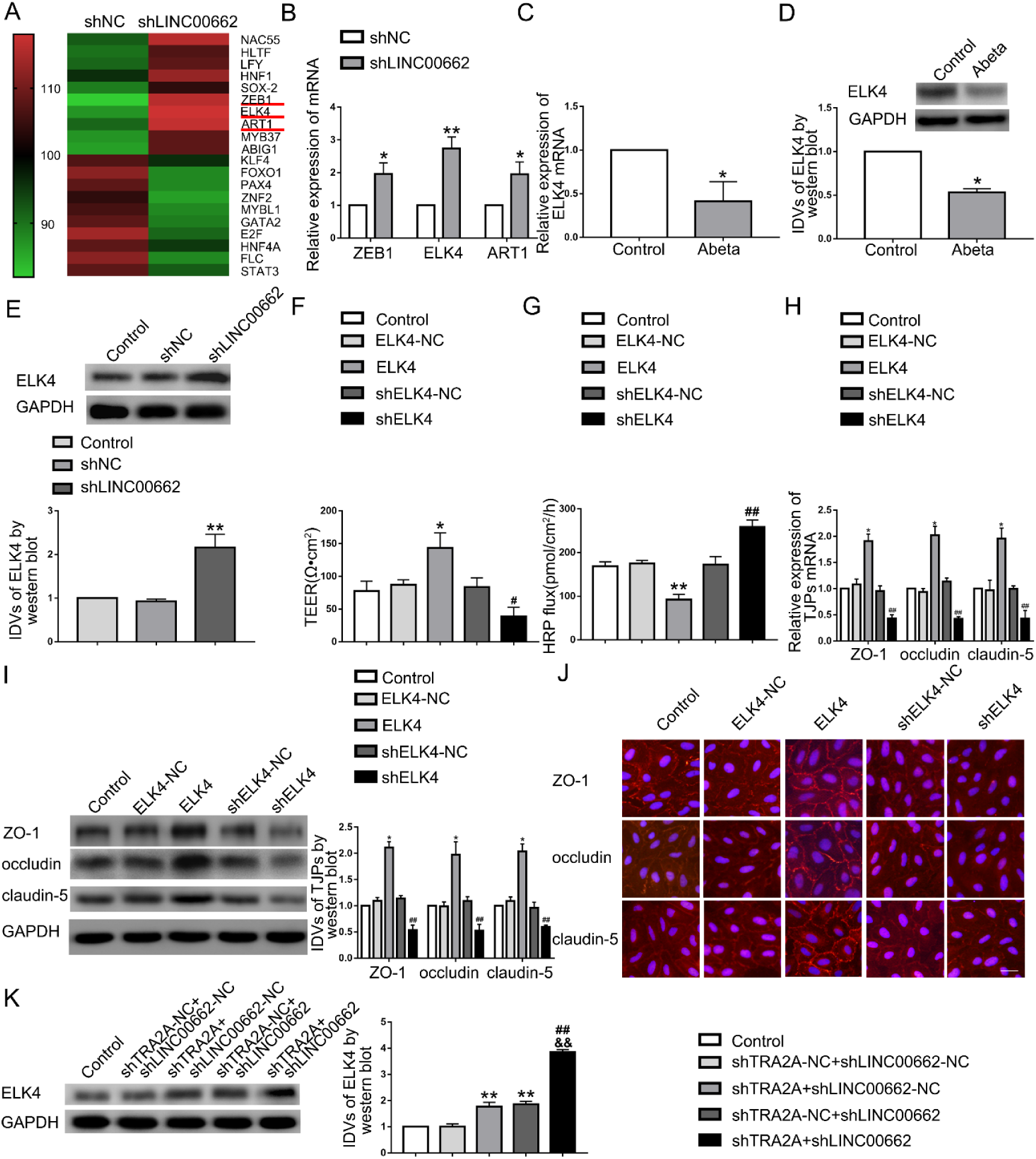
ELK4 expression in Aβ_1-42_-incubated ECs and ELK4 regulated BBB permeability in AD microenvironment. (A) RNA microarray analysis was performed in ECs treated with shLINC00662. Red indicates high relative expression and green indicates low relative expression. (B) Relative expression levels of ZEB1, ELK4, and ART1 determined by qRT-PCR. Data represent mean ± SD (n = 3, each). *\*\*P* < 0.01 *versus* shNC group. (C) Effects of LINC00662 on ELK4 expression level in Aβ_1-42_-incubated ECs by western blot. *\*\*P* < 0.01 *versus* control group. (D) Relative mRNA expression of ELK4 in ECs pre-incubated with Aβ_1-42_ by qRT-PCR. Data are presented as mean ± SD (n = 4, each). *\*P* < 0.05 *versus* control group. (E) Relative ELK4 protein levels in ECs pre-incubated with Aβ_1-42_ by western blot. *\*P* < 0.05 *versus* control group. (F and G) Effects of ELK4 on TEER values (F) and HRP flux (G). Data are presented as mean ± SD (n = 3, each). *\*P* < 0.05 *versus* ELK4-NC group. *\*\*P* < 0.01 *versus* ELK4-NC group. ^#^*P* < 0.05 *versus* shELK4-NC group. ^##^*P* < 0.01 *versus* shELK4-NC group. (H) Effects of ELK4 on ZO-1, occludin, and claudin-5 expression levels determined by qRT-PCR. Data are presented as mean ± SD (n = 3, each). *\*P* < 0.05 *versus* ELK4-NC group. ^##^*P* < 0.01 *versus* shELK4-NC group. (I) Effects of ELK4 on ZO-1, occludin, and claudin-5 expression levels determined by western blot. Data are presented as mean ± SD (n = 3, each). *\*P* < 0.05 *versus* ELK4-NC group. ^##^*P* < 0.01 *versus* shELK4-NC group. (J) Effects of ELK4 on ZO-1, occludin, and claudin-5 expression levels and distribution determined by immunofluorescence staining (n = 3, each). ZO-1, occludin, and claudin-5 (red) were labeled with secondary antibody against anti-ZO-1, anti-occludin, and anti-claudin-5 antibody, respectively, and nuclei (blue) were labeled with DAPI. Scale bar represents 30 μm. (K) ELK4 expression regulated by TRA2A and LINC00662 knockdown; data are presented as mean ± SD (n = 3, each).*\*\*P* < 0.01 *versus* shTRA2A-NC + shLINC00662-NC group, ^##^*P* < 0.01 *versus* shTRA2A + shLINC00662-NC, *^&&^P* < 0.01 *versus* shTRA2A-NC + shLINC00662 group.

### LINC00662 down-regulated the expression of ELK4 through SMD pathway, downregulated the expression of ZO-1, occludin and claudin-5, and increased the permeability of BBB in AD microenvironment

By quiring the bioinformatics database RepeatMasker and IntaRNA, we found that LINC00662 has the putative binding site with the ELK4 3’UTR (Figure. S-E-G). Therefore, we hypothesized that LINC00662 exhibits a negative regulatory effect on ELK4 *via* SMD pathway. The interaction between LINC00662, ELK4 and STAU1 was detected by dual-luciferase reporter assays, RNA pull down experiments and RIP. We used dual-luciferase reporter assays to confirm the predicted binding site. The relative luciferase activity in ELK4-3’UTR-Wt+LINC00662 group was remarkably lower than ELK4-3’UTR-Wt+LINC00662-NC group, while there was no significant difference between ELK4-3’UTR-Mut+LINC00662 and ELK4-3’UTR-Mut+LINC00662-NC group (Figure.5-A). RNA pull down experiments also confirmed that LINC00662 bind directly to ELK4 and STAU1, respectively (Figure. 5-B,D). Furthermore, as shown in Figure 5-C,E, RIP confirmed that LINC00662 and ELK4 mRNA bind to STAU1 directly respectively. The relative enrichment of LINC00662 and ELK4 mRNA in anti-STAU1 group was higher than anti-IgG group respectively. In order to confirm that STAU1 participates in the interaction between LINC00662 and ELK4, Aβ_1-42_-incubated ECs was co-tansfected with shSTAU1 and shLINC00662 plasmid. qRT-PCR demonstrated that nacent LINC00662 levels in control group, shSTAU1-NC+shLINC00662-NC group, shSTAU1+shLINC00662-NC group, shSTAU1-NC+shLINC00662 group and shSTAU1+shLINC00662 group are not statistically significant (Figure. 5-F). The half-life of ELK4 mRNA was significantly increased in shLINC00662+shSTAU1 group (Figure. 5-G). ShSTAU1+shLINC00662-NC and shSTAU1-NC+shLINC00662 groups exhibited higher ELK4 expression compared with shSTAU1-NC+shLINC00662-NC group respectively. The promotive effect was magnified in shSTAU1+shLINC00662 group (Figure. 5-H). To verify the effect of UPF1 on ELK4, ECs was transfected with shUPF1 plasmid. There were no statistical differences in nacent ELK4 mRNA between control group, shNC group and shUPF1 group. The half-life of ELK4 mRNA was significantly increased in shUPF1 group (Figure. S-J). ELK4 protein expression level increased significantly in shUPF1 group (Figure. S-K). As shown in Figure. 5-I, J, compared with shLINC00662-NC+ELK4-NC group, TEER in shLINC00662+ELK4 group increased and HRP flux decreased significantly. Co-knockdown of ELK4 and LINC00662 largely reversed the LINC00662 knockdown induced the increase in TEER and the impairment in HRP flux. As shown in Figure. 5-K, compared with shLINC00662-NC+ELK4-NC group, ZO-1, occludin, claudin-5 protein expression was increased in shLINC00662+ELK4 group. Moreover, co-knockdown of LINC00662 and ELK4 largely reversed the promotive effect of ZO-1, occludin, claudin-5 expression in Aβ_1-42_ incubated ECs caused by LINC00662 konckdown.

**Figure 5.**
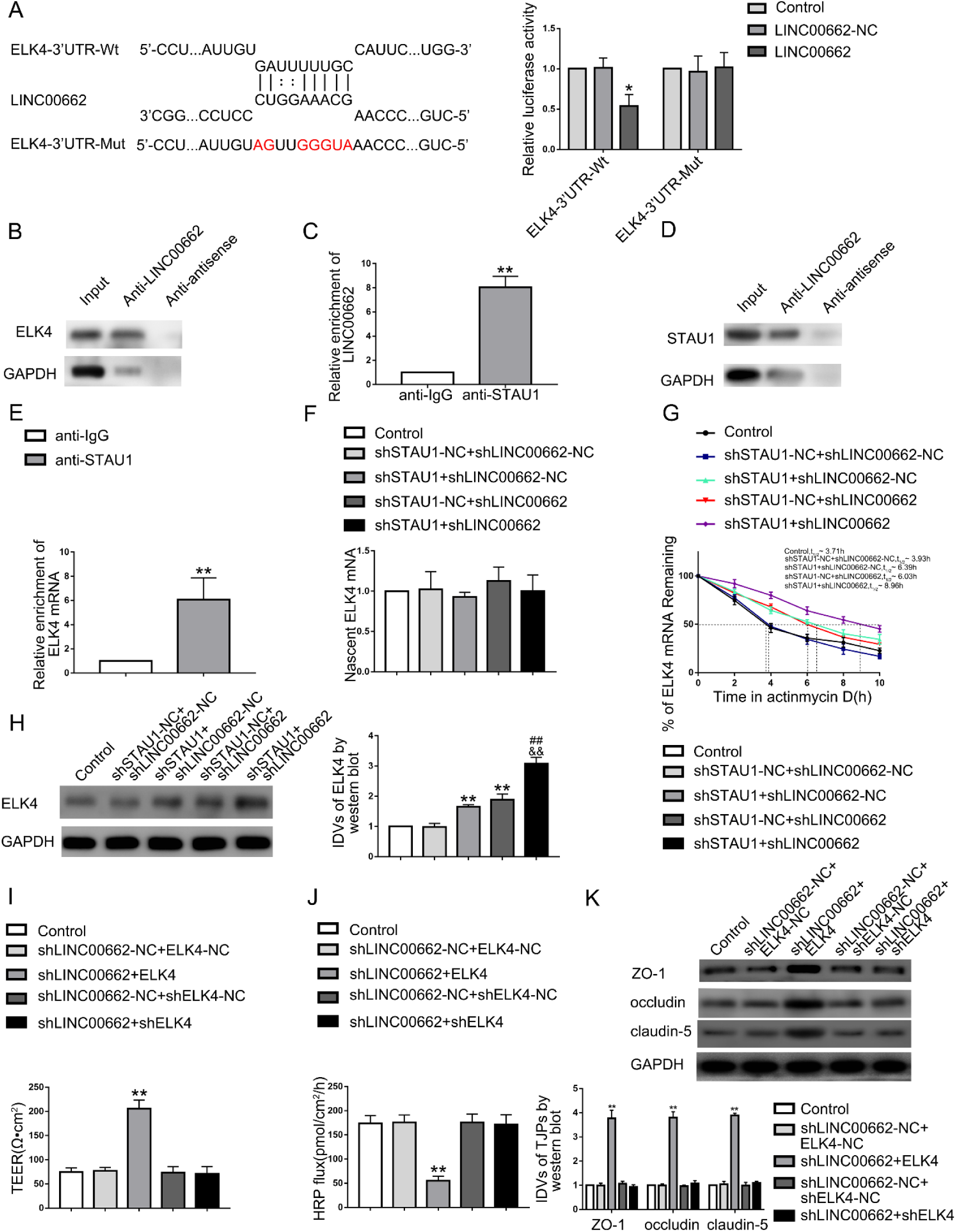
LINC00662 regulated BBB permeability by degrading ELK4 mRNA through SMD pathway. (A) The predicted LINC00662 binding site in ELK4 mRNA 3’UTR and results of dual-luciferase reporter assays. Data are presented as mean ± SD (n = 4, each group). *\*P* < 0.05 *versus* ELK4-3’UTR-Wt + LINC00662-NC group. (B) RNA-pull down confirmed the binding interaction between LINC00662 and ELK4. (C) RNA-IP confirmed the binding interaction between LINC00662 and STAU1. Relative enrichment was measured by qRT-PCR; data are presented as mean ± SD (n = 3, each). *\*\*P* < 0.01 *versus* anti-IgG group. (D) RNA pull-down indicated that LINC00662 bind to STAU1 directly. (E) RNA-IP confirmed the binding interaction between ELK4 and STAU1. Relative enrichment was measured by qRT-PCR; data are presented as mean ± SD (n = 3, each). *\*\*P* < 0.01 *versus* anti-IgG group. (F) The graph shows nascent ELK4 in Aβ_1-42_-incubated ECs treated with shLINC00662 and shSTAU1; data are presented as mean ± SD (n = 3, each). (G) Stability of ELK4 mRNA regulated by knockdown of LINC00662 and STAU1. (H) ELK4 expression regulated by STAU1 and LINC00662 knockdown; data are presented as mean ± SD (n = 3, each).*\*\*P* < 0.01 *versus* shSTAU1-NC + shLINC00662-NC group, ^##^*P* < 0.01 *versus* shSTAU1 + shLINC00662-NC, *^&&^P* < 0.01 *versus* shSTAU1-NC + shLINC00662 group. (I and J) TEER values (I) and HRP flux (J) to evaluate the effects of LINC00662 and ELK4 on BBB integrity. Data are presented as mean ± SD (n = 3, each). *\*\*P* < 0.01 *versus* shLINC00662-NC+ELK4-NC group. (K) Effects of LINC00662 and ELK4 on ZO-1, occludin, and claudin-5 expression levels determined by western blot. Data are presented as mean ± SD (n = 3, each). *\*\*P* < 0.01 *versus* shLINC00662-NC+ELK4-NC group.

### ELK4 bound to the promoters of ZO-1, occludin and claudin-5 and promoted transcription

To further verify the mechanisms of ELK4 affecting the permeability of BBB, we used bioinformatics software to query the relationship between ELK4 and tight junction-related proteins. We found that the ZO-1, occludin, claudin-5 promoter might harbor upstream putative binding sites for ELK4 according to JASPAR. Subsequent dual-luciferase reporter assays and ChIP assays were conducted to confirm any association between ELK4 and the mentioned promoters. As shown in Figure. 6-A-C, ZO-1, occludin, and claudin-5 promoter activities were remarkably increased after co-transfection with pEX3-ELK4. Subsequently, putative ELK4 binding sites witin the ZO-1, occludin, claudin-5 promoter were deleted successively. As shown in Figure 6-A, deletion of the -880 site and the -570 site caused reduction of the ZO-1 promoter activity, indicating that the -880 site and the -570 site of ZO-1 promoter contains high promoter activity. Whereas, after the deletion of the -381 site, the promoter activity didn’t change significantly. As shown in Figure 6-B, following deletion of the -531 site, the occludin promoter activity diminished. As shown in Figure 6-C, deletion of a -153 site produced reversion of promoter activity increase, and so dose deletion of a -81 site. Consistent with the dual-luciferase reporter assays, ChIP assays show that ELK4 bind with the ZO-1, occludin, and claudin-5 promoters on the same binding sites. As shown in Figure. 6-D-F, the specific primers for the -880, -570, -381 site regions of the ZO-1 promoter, -531 site region of the occludin promoter and -327, -153, -81 site regions of the claudin-5 promoter were used. In ELK4 immunoprecipitates, PCR constructs were observed using primers specific mentioned above and the 3000 bp upstream regions were used as negative control. In general, the results show that ELK4 transcriptionally promots ZO-1, occludin, and claudin-5 by binding with the gene’s promoters.

**Figure 6.**
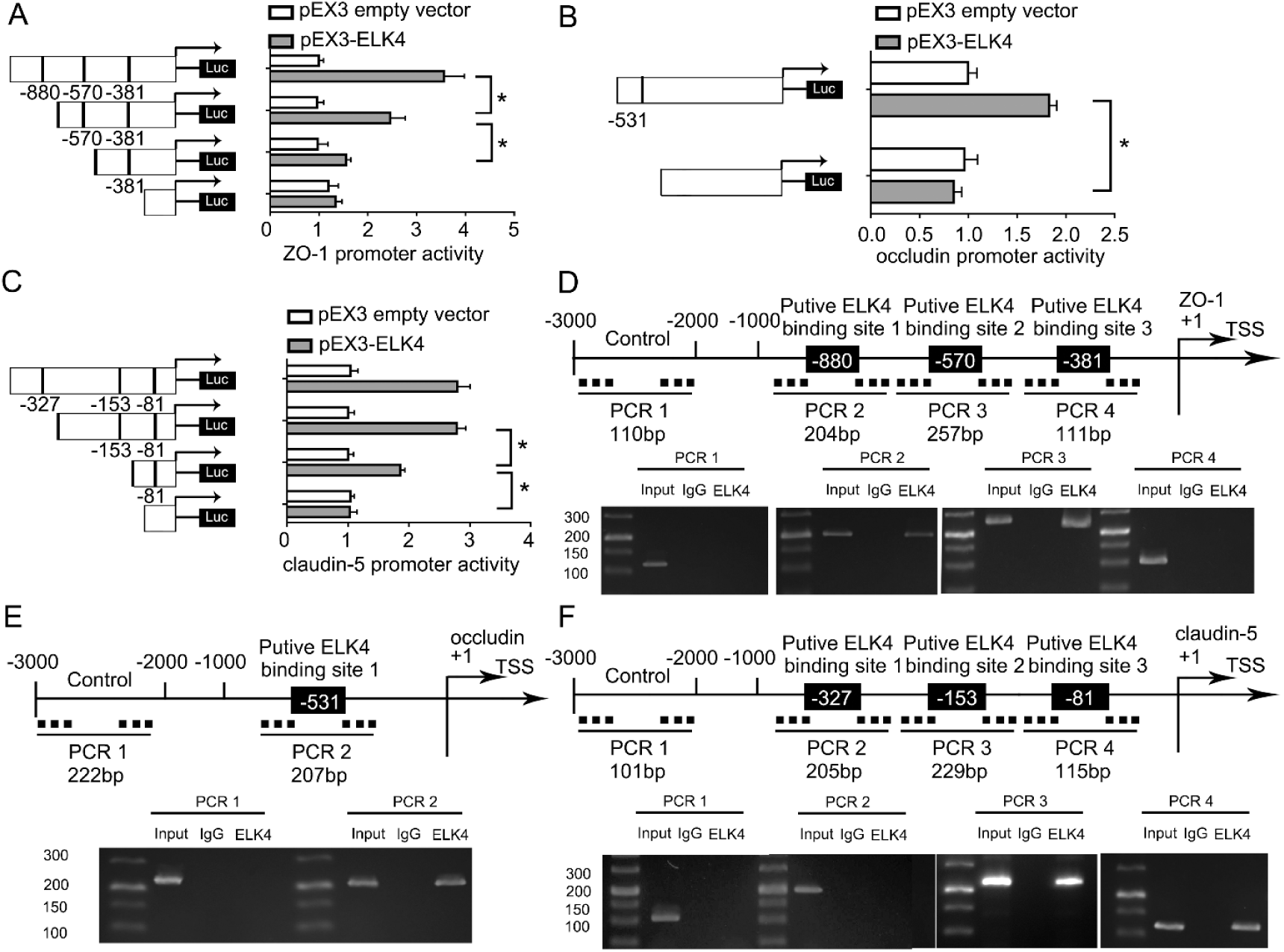
ELK4 increased the Promoter Activity of ZO-1, occludin, and claudin-5. (A-C) Schematic depiction of the different reporter plasmids and relative luciferase activity: ZO-1 (A), occludin (B), and claudin-5 (C) are shown. The Y-bar shows the deletion positions on the promoter fragments. The X-bar shows the reporter vector activity after normalization with the cotransfected reference vector (pRL-TK), and relative to the activity of the pEX3 empty vector, and the activity was set to 1. Data represent mean ± SD (n = 5, each). *\*P* < 0.05. (D-F) ELK4 bound to the promoters of ZO-1 (D), occludin (E), claudin-5 (F) in Aβ_1-42_-incubated ECs. Transcription start site (TSS) was designated as +1. Putative ELK4 binding sites are illustrated. Immunoprecipitated DNA was amplified by PCR. Normal rabbit IgG was used as a negative control.

## Discussion

Our study demonstrated for the first time that RBP-TRA2A and lncRNA-LINC00662 were highly expressed in Aβ_1-42_ incubated ECs, and knockdown of TRA2A or LINC00662 could significantly reduce BBB permeability in AD microenvironment. TRA2A knockdown reduced the stability of LINC00662 and down-regulated its expression; down-regulated LINC00662 attenuated the degradation of ELK4 mRNA through the SMD pathway, thereby increasing the expression of ELK4. Moreover, restoration of ELK4 decreased BBB permeability *via* transcriptional promotion of ZO-1, occludin, and claudin-5. The TRA2A/LINC00662/ELK4 axis is presented in Figure. 7 schematically.

**Figure. 7.**
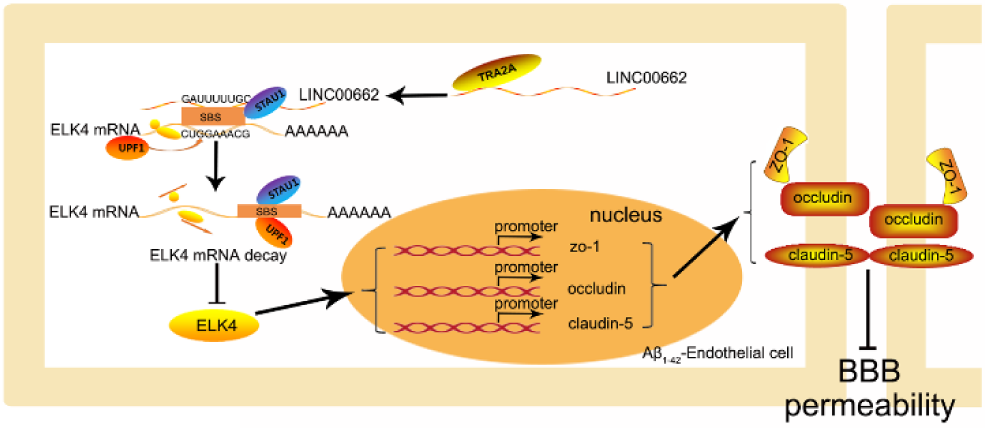
The schematic illusion of interactions between TRA2A, LINC00662 and ELK4 in Aβ_1-42_-incubated ECs.

BBB is important in maintaining the stability of physiological environment and brain functions of the central nervous system. BBB breakdown contributes to the onset and progression of AD^[26, 27]^. It has been found that BBB dysfunction causes diminishment of Aβ clearance, leading to Aβ deposition^[28–30]^. Transcellular and paracellular pathway are involved in the regulation of BBB permeability^[31]^. Tight junction-related proteins between ECs is important in the regulation of the paracellular pathway. Abnormal expression and distribution of tight junction-related proteins can affect BBB permeability ^[32]^. It has been reported that Aβ binds to receptor for advanced glycation endproducts (RAGE) and further induce the production of reactive oxygen species (ROS). The toxic effects on cerebral microvascular ECs leads to loss of tight junction-related proteins and disruption of BBB integrity^[33]^. Granulocyte-macrophage colony-stimulating factor (GM-CSF) decreases ZO-1 and claudin-5 expression by transcription modulation and the ubiquitination pathway, respectively. High GM-CSF levels in AD patients brain parenchyma and cerebrospinal fluid lead to blood-brain barrier open^[34]^. It has also been reported that BBB disruption due to lower expression and disassembly of tight junction-related proteins happens in age-associated brain changes and other neurodegenerative diseases^[35, 36]^.

Numerous reports have suggested that RBPs play an important regulatory role in neurodegenerative diseases. It has been suggested neuron-specific RBPs regulate several different neurological processes in AD. nELAVL proteins abundant in neuron-specific RBPs and bind to non-coding Y RNA forming nELAVL/Y RNA complex in AD. The complex results in nELAVL isolation leading to nELAVL sequestration, nELAVL target binding redistribution, and neuronal RNA splicing alteration^[37]^. Mutations in RBP fused in sarcoma (FUS) gene contribute to familial Amyotrophic lateral sclerosis (ALS) cases. Mutant FUS reduces protein synthesis by disrupting the nonsense-mediated decay (NMD) pathway, which may lead to motor neuron death^[38]^. TRA2A was confirmed to be induced in Aβ_1-42_-incubated ECs. Knockdown of TRA2A attenuated BBB permeability in AD microenvironment by increasing the expression of ZO-1, occludin and claudin-5. RBPs have played an pivotal role in regulating vascular endothelial function. RBP QKI is highly expressed in ECs and promotes the mRNA of β-catenin and VE-cadherin translation by binding with them, which contributes to maintenance of endothelial barrier function^[39]^. RBPs MOV10 and FUS regulate vascular endothelial function by binding to Circular RNAs (circRNAs)^[40, 41]^.

LncRNAs is signigicant in brain development and neurodegenerative diseases including epigenetic regulation, transcriptional regulation, post-transcriptional regulation, translational regulation^[42, 43]^. In Huntington’s disease, neurodegeneration is related with lncRNAs expression changes, for many of the changes cause the alteration of epigenetic gene regulation in neurons^[44]^. A novel lncRNA (named 17A) is enriched in AD patients’s brain tissues and embeds in the human G-protein-coupled receptor 51 gene (GPR51, GABA B2 receptor). 17A promotes abolishment of GABA B2 intracellular signaling in SHSY5Y neuroblastoma cells and enhances the secretion of Aβ. Thus 17A contributes to the occurrence and development of AD^[45]^. Highly expression of lncRNA NDM29 increased Aβ through the promotion of Aβ molecular processing and secretion in AD^[46]^. In our study, LINC00662 was highly expressed in Aβ_1-42_-incubated ECs. Knockdown of LINC00662 decreased BBB permeability in AD microenvironment by increasing the expression of tight junction-related proteins. Accumulated evidences have indicated that lncRNAs regulate vascular endothelial function. LncRNA-CCL2 positively regulates CCL2 mRNA levels in ECs, which may contribute to endothelial inflammatory changes^[47]^. LncRNA SENCR promotes EC adherens junction integrity through physical association with CKAP4, thereby stabilizing cell membrane^[48]^. LINC00094 affects blood-brain barrier permeability in AD microenvironment by regulating the expression of tight junction-related proteins^[32]^.

Recently, intense studies have indicated that the interaction between RNA-binding proteins and non-coding RNAs regulates vascular endothelial function. We found that TRA2A can increase the stability of LINC00662 and reduce BBB permeability in AD microenvironment. One of the mechanisms is that RBPs alter the stability of non-coding RNAs and regulate vascular endothelial function. For example, Wu *et al*^[49]^ have reported that RBP KHDRBS3 binds to circRNA DENND4C, increases the circRNA’s stability and upregulates the expression of tight junction-related proteins, thereby reducing the permeability of blood tumor barrier. Li *et al*^[15]^ have reported that RBP ZRANB2 binds to the lncRNA and increases the lncRNA’s stability, thereby affecting the functions of vascular ECs. A similar report has been made in AD that RBP HuD associates with and stabilizes the lncRNA BACE1AS, which partly complements encoding β-site APP-cleaving enzyme 1 (BACE1) mRNA. HuD elevates the levels of APP, BACE1, BACE1AS, and promotes the production of APP and the cleavage of its amyloidogenic fragment, Aβ^[50]^.

ELK4 is a member of the Ets transcription factor family, which is a critical regulator of many neurodevelopmental events^[51]^. ELK4 can not only bind to gene promoter directly but also cooperate with other transcription factors to regulate target gene transcription^[19, 52]^. ELK4 upregulates Mcl-1 by binding to Mcl-1 promoter and improves its activity, thereby decreasing glioma cells sensitivity to apoptosis^[21]^. In HepG2 cells, ELK4 is a protein through which SIRT7 targets the G6PC promoter, regulates the transcriptional activity of G6PC under glucose deprivation, and participates in the regulation of gluconeogenesis^[53]^. ELK4 is enriched in promoter bindings of genes upregulated in peripheral blood mononuclear cells (PBMC) from non-small cell lung cancer (NSCLC) patients before tumor removal, indicating that transcriptional promotion function of ELK4 is involved in the interaction between NSCLC and immunologically important pathways^[54]^. In present study, dual-luciferase gene reporter assays and ChIP assays were used to demonstrate for the first time that ELK4 could transcriptionally promoted ZO-1, occludin, claudin-5 and reducing BBB permeability in AD microenvironment.

Current studies have shown that SMD pathway has an important significance in neurodevelopment and neurological diseases. For example, SMD pathway is promoted by KLF4 through increasing Stau1 recruitment to the 3’UTR of neurogenesis-associated mRNAs. The molecular mechanism underlying stability of neurogenesis-associated mRNAs affects neurogenesis and self-renewal in mice^[55]^. In addition, in mouse 3T3-L1 cells, PNRC2 interacts with STAU1 and UPF1 to reduce the stability of KLF2 mRNA and accelerates adipogenesis through SMD pathway^[24]^. We found that knockdown of LINC00662, STAU1 and UPF1 could prolong the half-life of ELK4 improving the stability of ELK4 mRNA and increase the expression of ELK4. Our study proved, for the first time, that LINC00662 promoted ELK4 degradation *via* SMD pathway in ECs, thus increasing BBB permeability in AD microenvironment. Earlier studies have shown that lncRNAs regulate the function of vascular ECs and tumor cells by SMD pathway^[56]^. For example, SNHG20 reduces FOX1 mRNA stability through SMD pathway, thereby regulating vascular ECs^[15]^. In the SGC7901 and BGC823 cell lines, lncRNA TINCR impairment the stability of KLF12 mRNA *via* SMD pathway, affecting the proliferation and apoptosis of gastric cancer cells^[57]^.

In conclusion, we studied TRA2A and LINC00662 expression increased and ELK4 expression impaired in Aβ_1-42_-incubated ECs. TRA2A bound to LINC00662, and LIN00662 acted on ELK4 *via* SMD pathway. Knockdown of TRA2A reduced the stability of LINC00662, and reduced the degradation of ELK4 mRNA by SMD pathway subsequently. ELK4 promoted transcription by binding to the promoters of ZO-1, occludin and claudin-5 and further reduced the permeability of blood-brain barrier in AD microenvironment. More significantly, TRA2A, LINC00662 and ELK4 may have a significant therapeutic potential in AD treatment. Our results provide a new mechanistic insight into the pathogenesis of AD and identify novel targets for its treatment.

## Materials and methods

### Cell cultures

The human cerebral microvascular endothelial cell line hCMEC/D3 (ECs) were provided by Dr. Couraud (Institut Cochin, Paris, France). Human brain vascular pericytes (HBVP) and normal human astrocytes (NHA) were purchased from the Sciencell Research Laboratories (Carlsbad, CA, USA). NHA and HBVP performed in the research were limited with passage below 12. ECs (passage 28-32), HBVP (passage 10-12) and NHA (passage 10-12) were cultured as described previously^[58]^. Human embryonic kidney 293 (HEK293T) cells were purchased from Shanghai Institutes for Biological Sciences Cell Resource Center and the cell culture has been previously detailed^[59]^. All cells were maintained at 37°C, 5% CO2, in a humidified atmosphere. Aβ_1-42_ was manufactured by Sigma-Aldrich (St. Louis, MO, USA). Aβ_1-42_ was first dissolved in dry DMSO (2 mmol/L) and stored at −20°C. 2 mmol/L Aβ_1-42_ in DMSO was diluted into 200 μmol/L in cold Opti-MEM media and incubated at 4°C for 24 hours. Cells were pre-incubated with Aβ_1-42_ at a concentration of 5μmol/L for 48 hours following Liu *et al* ^[58]^.

### *In vitro* BBB model establishment

*In vitro* co-culturing BBB models were established as described previousely^[58]^. First, after pericytes were cultured (2 × 10^5^ cells/cm^2^) on the lower chamber of Transwell inserts (0.4 μm pore size; Corning, NY, USA) overnight, hCMEC/D3 cells were subsequently placed on the upper chambers of Transwell inserts. NHA (2×10^5^ cells/cm2) were seeded onto the 6-well culture plate and cultured for 2 days before adding ECs inserts.

### Real-time PCR assays

Nanodrop Spectrophotometer (ND-100, Thermo Scientific, Waltham, MA) was applied to determine the RNA concentration and quality. The expression levels of TRA2A (NM_001282757), LINC00662 (NR_027301) and ELK4 (NM_001973.4) were detected by One-Step SYBR PrimeScript RT-PCR Kit (Perfect Real Time; Takara Bio, Inc., Kusatsu, Japan). Relative expression values were calculated using the relative quantification (2^−△△Ct^) method. Primers were shown in Table A1.

### Cell transfections

Silencing plasmid of LINC00662 was ligated into LV10 (U6/RFP&Puro) vector (GenePharma, Shanghai, China) to construct the shLINC00662 plasmid. Short-hairpin RNA directed against human TRA2A gene, STAU1 gene, UPF1 gene and ELK4 gene was ligated into the pGPU6/GFP/Neo vector (GenePharma) to construct shTRA2A, shSTAU1, shUPF1, and shELK4 plasmid, respectively. The human ELK4 gene coding sequence was ligated into pIRES2 vector (GenScript, Piscataway, NJ, USA) to construct the ELK4 overexpression plasmid. The respective no-targeting sequences were used as NCs. The ECs were stable transfected *via* LTX and Plus reagent (Life Technologies, Carlsbad, CA, USA). The stable transfected cells were selected using G418 (0.4 mg/ml) (Sigma-Aldrich, St. Louis, MO, USA) or puromycin (1 μg/ml) (Sigma-Aldrich). Sequences of shTRA2A, shLINC00662, shELK4, shSTAU1, shUPF1 were shown in Table A2. The silencing and over-expressions efficiency were measured by qRT-PCR. The transfected efficiency of TRA2A, LINC00662 and ELK4 were shown in Figure. S-A-C, H, I. For co-transfection of shTRA2A and shLINC00662 (shTRA2A+shLINC00662), shLINC00662 was transiently transfected into Aβ_1-42_-incubated ECs, which stably transfected shTRA2A with lipofectamine 3000 reagent. After 48 hours, the transiently transfected cells were obtained. Other co-transfected ECs were established in the same way.

### Transendothelial electric resistance (TEER) assays

A millicell-ERS apparatus (Millipore, Billerica, MA, USA) was applied to perform TEER assays after *in vitro* BBB models constructed. Each measurement was placed in room temperature for 30 minutes, and TEER values was recorded. TEER values were measured after erchanging the medium. Background electrica resistances were subtracted before the final resistances were calculated. TEER values (Ω·cm^2^) is electric resistance multiplied by the effective surface area of the transwell insert.

### Horseradish peroxidase (HRP) flux assays

*In vitro* BBB models were constructed and 1ml of serum-free EBM-2 medium containing 10 μg/mL HRP (0.5 mmol/L, Sigma-Aldrich) culture medium was added into the upper champer of the transwell system. 1 hour later, 5 μL of culture medium in the lower chamber was collected and the HRP content of the samples was detected by tetramethylbenzidine colorimetry approach. The final HRP value was expressed as pmol/cm^2^/h.

### Western blot assays

The cell lysates were extracted from ECs. Total proteins were extacted with RIPA buffer (Beyotime Institute of Biotechnology, Jiangsu, China) supplemented with protease inhibitors (10 mg/mL aprotinin, 10 mg/mL phenyl-methylsulfonyl fluoride[PMSF], and 50 mM sodium orthovanadate) and centrifuged at 17,000×g for 30 minutes at 4°C. Equal amounts of proteins were further separated using SDS-PAGE and then transferred to polyvinylidene fluoride (PVDF) membranes (Millipore, Shanghai, China). Membranes were blocked to avoid non-specific bindings in Tris-buffered saline-Tween (TBST) containing 5% fat-free milk for 2 hours and subsequently incubated with primary antibodies (shown in Table A4) at 4°C overnight. After three washes with TBST, membranes were incubated with the corresponding secondary antibody at a 1:10000 dilution at room temperature for 2 hours. Immunoblots were visualized by enhanced chemiluminescence (ECL kit, Santa Cruz Biotechnology) after washes. All the protein bands were scanned by Chem Imager 5500 V2.03 software and the integrated density values (IDVs) were calculated utilizing FluorChem 2.0 software.

### Immunofluorescence assays

After fixed by 4% paraformaldehyde for 20 minutes, the cells permeated in phosphatebuffered saline (PBS) containing 0.2% Triton X-100 for 10 min. Next, cells were blocked by 5% bovine serum album (BSA) in PBS for 2 hours at room temperature, and incubated with primary antibodies (anti-ZO-1, 1:50, Life Technologies; anti-occludin, 1:50, Life Technologies; anti-claudin-5, 1:50, Life Technologies;) respectively at 4°C overnight. After three washes with PBS, cells were incubated with fluorophore-conjugated secondary antibodies for 2 hours. DAPI were applied to observe cell nuclei. The staining was oberservrd by immunofluorescence microscope (Olympus, Tokyo, Japan).

### Chromatin immunoprecipitation assays

Chromatin immunoprecipitation (ChIP) kit (Cell Signaling Technology, Danvers, MA, USA) was used for ChIP assays following the manufacture’s description. Briefly, cells were crosslinked with 1% formaldehyde and collected in lysis buffer. Chromatin was then digested with Micrococcal Nuclease. Immunoprecipitates were incubated with 3 μg anti-ELK4 antibody (Santa Cruz, USA) or normal rabbit IgG and incubated with Protein G agarose beads at 4°C overnight with gentle shaking while 2% lysates were used as input reference. DNA crosslink was reversed with 5M NaCl and proteinase K and purified. Immunoprecipitation DNA was amplified by PCR using their spcific primers. In each PCR reaction, the corresponding inputs were taken in parallel for PCR validation. Primers used for ChIP PCR are shown in Table A3.

### Human lncRNA and RNA microarrays

LncRNA and RNA analysis, sample preparation, and microarray hybridization were completed by Kangchen Bio-tech (Shanghai, China).

### Reporter vector construction and dual-luciferase reporter assays

The putative LINC00662 binding regions within the ELK4 gene were amplified by PCR and cloned into downstream of pmirGLO dual-luciferase vector (Promega, Madison, WI, USA) to form the wide-type plasmid (ELK4-3’UTR-Wt) (GenePharma). Similarly, the binding sequences were mutated as indicated to form the mutant-type plasmid (ELK4-3’UTR-Mut). The pmirGLO vector constructed with either 3’-UTR fragments or mutation of 3’-UTR fragments, and LINC00662 or LINC00662-NC were cotransfected into HEK293T cells in 24-well plates using Lipofectamine 3000. ECs were obtained for analysis by luciferase assay using the Dual-Luciferase Reporter Assay System (Promega) 48 houes after transfection. The relative luciferase activity was expressed as the ratio of firefly luciferase activity to renilla luciferase activity.

Human genomic DNA was used to amplify different promoter fragments, subcloned into pGL3-Basic-Luciferase vector (Promega) containing a firefly luciferase reporter gene and verified by DNA sequencing. Human full-length ELK4 was constructed in pEX3 (pGCMV/MCS/Neo) plasmid vector (GenePharma). HEK293T cells were co-transfected with the pGL3 vector of ZO-1, occludin, and claudin-5 with either full-length promoter regions (or deleted promoter regions) and pEX3-ELK4 (or pEX3 empty vector) using Lipofectamine 3000. Relative luciferase activity was analyzed as described previously.

### RNA immunoprecipitation (RIP) assays

EZ-Magna RNA-binding protein immunoprecipitation kit (Millipore, USA) was performed according to the manufacture’s protocol. Whole cell lysate was incubated with human 5 μg human anti-Ago2 antibody, or NC normal mouse IgG. Furthermore, purified RNA was extracted and applied to qRT-PCR to demonatrate the presence of the binding targets.

### RNA pull-down assays

Biotin-labelled, full length LINC00662, or antisense RNA was prepared with the Biotin RNA Labeling Mix (GenePharma, Shanghai, China) and transfected into ECs. Biotinylated RNAs were treated with RNase-free DNase I and purified. RNA-protein complexes were isolated by streptavidin agarose beads (Invitrogen, Shanghai, China) and washed three times. The retrieved proteins were detected using a standard western blotting technique with GAPDH as the control.

### Nascent RNA capture

Nascent RNAs was detected using Click-iT® Nascent RNA Capture Kit (Thermo Fisher Scientific,USA) according to the manufacture’s protocol. Briefly, nascent RNAs were marked with 0.2mM 5-ethymyl uridine (EU) and the EU-nascent RNA was captured on magnetic beads for subsequent qRT-PCR.

### mRNA stability assays

In order to inhibit the nascent RNA synthesis, 5 ug/ml actinomycin D (ActD, NobleRyder, China) was added into ECs culture medium. Total RNA was extracted at 0, 2, 4, 6, 8, 10h and its concentrations were easured by qRT-PCR. The half-life of RNA was determined by its level at certain point of time compared with time zero.

### Satistical analysis

Statistical analysis was performed with GraphPad Prism v7.00 (GraphPad Software, La Jolla, CA, USA) software. Data was described as mean ± standard deviation (SD). All differences were analyzed by SPSS 18.0 statistical software with the Student’s t-test (two tailed) or one-way ANOVA. *P*-values below 0.05 were considered significant.

## Authors’ Contributions

YXX contributed to the experiment design and implementation, manuscript draft, and data analysis. QSL contributed to the experiment implementation and data analysis. YHL conceived or designed the experiments. QSL, LZ, XBL, and JZ performed the experiments. LZ, XLR, SC, HC and ZL analyzed the data. QSL conceived or designed the experiments, performed the experiments, and wrote the manuscript. All authors read and approved the final manuscript.

---

**Supplementary Figure.**
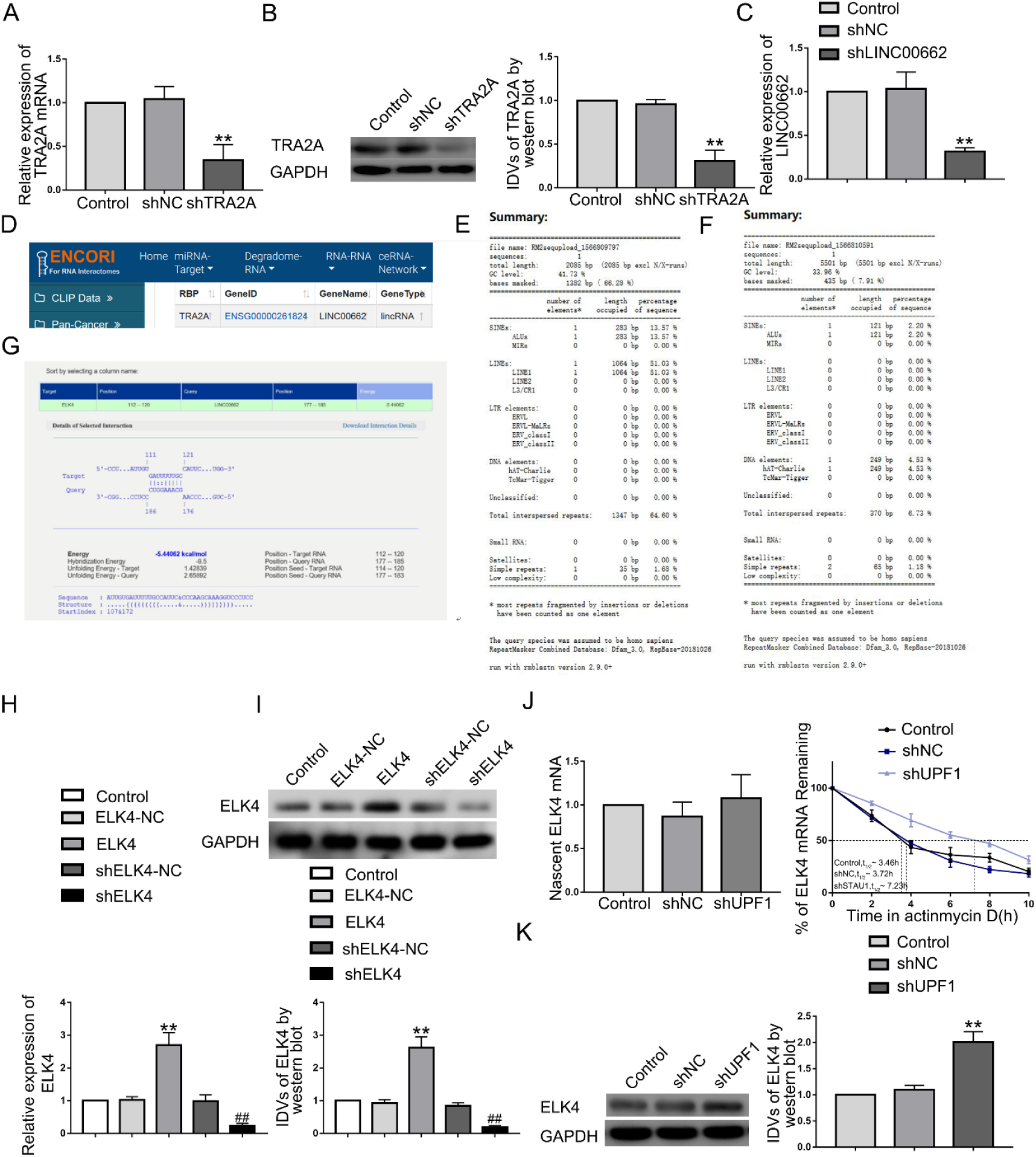
(A) Relative expression of TRA2A evaluated using qRT-PCR in the TRA2A knockdown Aβ_1-42_-incubated ECs; data are presented as mean ± SD (n = 3). *\*\*P* < 0.01 *versus* sh-NC group. (B) Relative expression of TRA2A evaluated by western blot in the TRA2A knockdown Aβ_1-42_-incubated ECs; data are presented as mean ± SD (n = 3). *\*\*P* < 0.01 *versus* sh-NC group. (C) Relative expression of LINC00662 evaluated using qRT-PCR in the LINC00662 knockdown Aβ_1-42_-incubated ECs; data are presented as mean ± SD (n = 3). *\*\*P* < 0.01 *versus* sh-NC group. (D) The predicted combination of TRA2A and LINC00662 by starBase. (E and F) The Alu elements in LINC00662 and ELK4 3’UTR by RepeatMasker. (G) The predicted LINC00662 binding site in ELK4 mRNA 3’UTR by IntaRNA. (H) Relative expression of ELK4 evaluated using qRT-PCR in ELK4 knockdown and overexpression Aβ_1-42_-incubated ECs; data are presented as mean ± SD (n = 3). *\*\*P* < 0.01 *versus* ELK4-NC group. ^##^*P* < 0.01 *versus* shELK4-NC. (I) Relative expression of ELK4 was evaluated by western blot in ELK4 knockdown and overexpression Aβ_1-42_-incubated ECs; data are presented as mean ± SD (n = 3). *\*\*P* < 0.01 *versus* ELK4-NC group. ^##^*P* < 0.01 *versus* shELK4-NC. (J) Stability of ELK4 mRNA treatde with shUPF1; data are presented as mean ± SD (n = 3, each group). (K) Effects of UPF1 on ELK4 expression; data are presented as mean ± SD (n = 3, each group). *\*\*P* < 0.01 *versus* shNC group.

